# Seroprevalence and cross-reactivity of neutralizing antibodies to six types of adenoviruses

**DOI:** 10.1101/2024.12.26.630365

**Authors:** O.V. Zubkova, T.A. Ozharovskaia, O. Popova, I.V. Vavilova, D.I. Zrelkin, P.P. Goldovskaya, D.V. Voronina, I.V. Dolzhikova, D.M. Grousova, D.V. Shcheblyakov, D.Y. Logunov, A.L. Gintsburg

**Affiliations:** Federal State Budgetary Institution “National Research Center for Epidemiology and Microbiology named after Honorary Academician N.F. Gamaleya”, Ministry of Health, Russian Federation

**Keywords:** adenovirus, vector vaccine, cross-neutralization, seroprevalence

## Abstract

**Background:** The use of adenoviral vector vaccines is an effective way to induce an antiviral immune response. Most vaccines used worldwide are based on human adenovirus type 5 (Ad5). Pre-existing natural immunity to Ad5 is a limitation to the frequent use of Ad5-based vectors in gene therapy and vaccination. In the present study, we assessed the seroprevalence and cross-reactivity of neutralizing antibodies to six adenovirus types: Ad1, Ad2, Ad5, Ad19, Ad26, and Ad37.

**Methods:** The study used immunological and virological research methods. Adenovirus vectors encoding the enhanced green fluorescent protein (EGFP) were developed by the N.F. Gamaleya National Research Center for Epidemiology and Microbiology. For seroprevalence and cross-neutralation study we used a panel of sera from 101 volunteers.

**Results:** The conducted correlation analysis showed a strong relationship between seropositivity and the geometric mean titer of neutralizing antibodies to adenovirus. An increase in antibody titers indicates the formation of a humoral immune response and population immunity. We determined that there is no cross-reactivity between NtAbs to Ad1, Ad2, Ad5 and Ad26.

**Conclusions:** Based on the presented data, we assume that Ad1 or Ad2 should be considered the most promising alternative candidates for the development of vaccine vectors.

## Introduction

To date, adenoviruses remain an important object in various medical studies. Recombinant adenoviruses play the key role in the development of vector vaccines, gene therapy, oncolytic therapy and genetic engineering [1]. The use of adenoviral vector vaccines is an effective way to induce an antiviral immune response. Numerous studies of vaccines based on recombinant human adenovirus type 5 (Ad5) have shown the ability to form protective adaptive immunity after vaccination [2-8]. However, according to the results of clinical studies, pre-existing natural immunity to adenoviruses in the human population limits the use of adenoviral vectors [9]. Human adenoviruses types 3, 5 and 7 (Ad3, Ad5 and Ad7) are the cause of acute respiratory infections development [10,11]. Thus, serological studies in Guangzhou (China) have shown a high level of neutralizing antibodies against these types of adenoviruses: 73.1% of the adult population are seropositive to Ad5; 20-85% - to Ad3 and Ad7 [12-16]. This fact is associated with a significant suppression of the immunogenicity of the Ad5-based vaccine against HIV-1 [8, 17-19]. Therefore, at present, the urgent task is to study rare types of adenoviruses characterized by low seroprevalence in the human population [20-22]. Despite the large number of known types of adenoviruses, not all of them are promising for developing recombinant vectors. The developed adenovirus vectors must have high immunogenicity in the presence of pre-existing immunity to the most common adenoviruses (to Ad5 particularly).

One of the first alternative vectors with low seroprevalence was human adenovirus type 35 (Ad35) [23]. Recombinant vectors based on Ad35 showed efficient transduction of dendritic cells, providing a high level of delivery and presentation of antigen [24-27]. The next alternative adenovirus to be studied was human adenovirus type 26 (Ad26) [11, 23].

Adenoviral vector vaccines for the prevention of COVID-19 have been approved for clinical use and have shown good results [28]. The Sputnik-V vaccine, developed in the Gamaleya Research Center, includes two adenoviral vectors, Ad26 and Ad5, and is used for heterologous prime-boost immunization to induce robust immunity [29]. The Chinese COVID-19 vaccine Convidecia, produced by CanSinoBio, is based on Ad5. Clinical studies have shown that pre-existing natural immunity in the population affects the effectiveness of vaccination. After vaccination, antibodies against SARS-CoV-2 coronavirus in a group of volunteers with a high level of pre-existing immunity to Ad5 (antibody titer > 1:200) were 2 times lower than in the control group [5]. Oxford-AstraZeneca used a vector based on chimpanzee adenovirus type 68 (ChAd68) to produce the ChAdOx1 vaccine. Although antibodies to ChAd68 are relatively rare in the human population, high levels of neutralizing antibodies (NAbs) to the vector were detected in individuals immunized with the ChAdOx1 vaccine [30]. These facts should be taken into account when developing adenovirus-based vaccines.

In recent years, a significant number of studies have been conducted in healthy volunteers to identify human adenovirus (Ad) types with low seroprevalence [11, 14, 15]. Taken together, these studies have shown that the seroprevalence of neutralizing antibodies to different Ad types varies significantly both in terms of reactivity and across countries and even regions within countries [31]. In the present study, we assessed the seroprevalence and cross-reactivity of neutralizing antibodies to six adenovirus types: Ad1, Ad2, Ad5, Ad19, Ad26, and Ad37. To our knowledge, this is the first study to assess the correlation of NAbs levels between different Ad types.

## Materials and methods

### Cell lines

HEK293 cells were cultured in DMEM medium (Cytiva, Austria) containing 8% fetal bovine serum (Gibco, USA), 25,000 U penicillin and 25 mg streptomycin (Paneko, Russia) and in a humidified atmosphere with 5% CO_2_ at 37 °C.

### Adenoviral vectors

Ad1, Ad2, Ad5, Ad19, Ad26 and Ad37 vectors encoding the enhanced green fluorescent protein (EGFP) were obtained as described previously [32,33]. Viruses were propagated in HEK293 cells and purified using a standard cesium chloride gradient ultracentrifugation method [34]. The number of viral particles (vp/ml) was determined spectrophotometrically as described previously [35]. The number of infectious particles was determined using the tissue culture infectious dose 50% (TCID_50_) assay as described previously [36].

### Serum samples and ethical approval

Serum samples and serum dilutions were stored at -20 °C. This study was approved by the Biomedical Ethics Committee of the Gamaleya National Research Center for Epidemiology and Microbiology of the Russian Ministry of Health (Protocol No. 81 dated 05/11/2024).

### Determination of NtAb titer to Ad5 in blood sera

NtAb titer to Ad was determined in a microneutralization assay on a HEK293 cell line using Ad expressing the EGFP gene at a dose of 100 TCID_50_. To conduct the assay, blood sera were inactivated at 56 °C for 30 minutes. Ad and blood sera dilutions were prepared in serum-free DMEM medium. 50 μl of two-fold serum dilutions were added to the wells of a 96-well plate, starting with 1:25, then 50 μl of the virus-containing suspension were added, after which the plates were incubated for 1 hour at 37 °C. Each sample was titrated in triplicate. After the incubation period, 100 μl of the cell suspension were added to each well of the plate at a rate of 5-6 × 10^6^ cells per plate. The cells were incubated at 37 °C and 5% CO_2_ for 48-72 h. The results of the neutralization reaction were recorded using UV microscopy on an Olympus IX73 inverted microscope (Olympus; Japan) with a U-RFL-T fluorescence module (Olympus; Japan). The NtAb titer of the serum under study was taken as its highest dilution, which suppressed virus replication in HEK293 cells by 50% of the positive control.

### Statistical analysis

Statistical analysis was performed in GraphPad Prism 8.4.2 using one-way analysis of variance (ANOVA) on ranks to compare antibody levels among groups. Spearman’s correlation coefficient was used to determine the degree of correlation between NtAb titers to Ad. Correlation analysis of seroprevalence and geometric mean virus-neutralizing antibody titer was performed using Pearson’s method. P-values less than 0.05 was considered significant.

## Results

### Adenovirus seroprevalence

To test the possibility of using other types of adenoviruses as alternative options in the development of vector systems, we studied the titers of neutralizing antibodies to Ad1, Ad2, Ad5, Ad19, Ad26, and Ad37 in the blood serum of volunteers (Fig. 1). NtAbs were detected in a microneutralization assay using recombinant adenoviral vectors carrying EGFP as a reporter gene. We screened a panel of sera from 101 volunteers and found a minimal number of Ad37-seropositive sera. The proportion of Ad37-positive samples was 10.89% (Fig. 1A). The percentage of Ad19-positive samples was 24.75. Less than half of the samples (46.53%) were seropositive to Ad1. NtAbs to Ad2, Ad5, and Ad26 were detected in 80.2%, 62.38%, and 58.42% of the analyzed sera, respectively.

**Figure 1.**
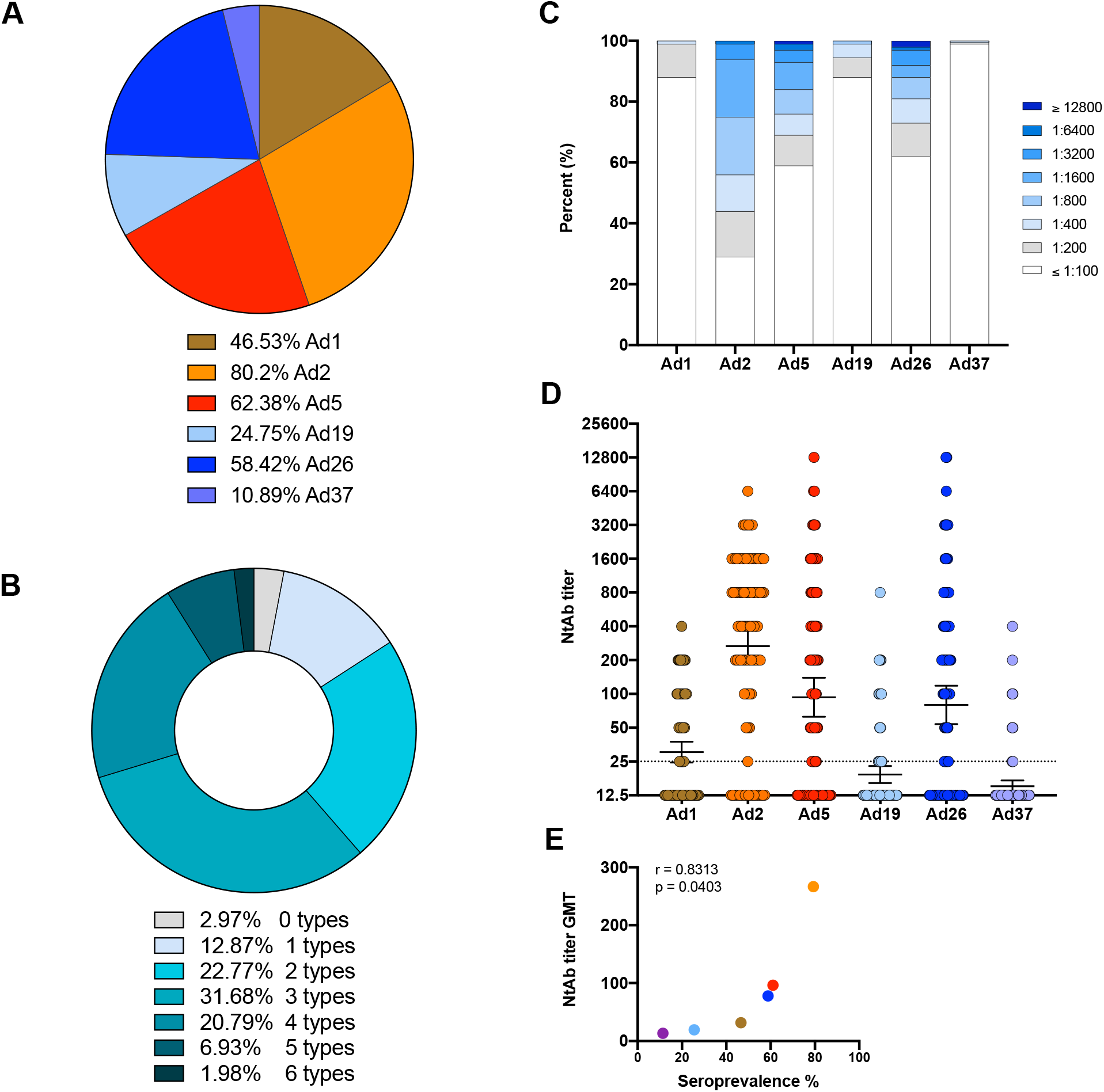
Seroprevalence of neutralizing antibodies to adenoviruses. (A) Structure of adenovirus seroprevalence. Serum samples from 101 adults from Russia were evaluated for Ad1-, Ad2-, Ad5-, Ad19-, Ad26-, Ad37-specific NtAbs. NtAb titers <25 were considered negative. (B) Percentage of serum samples that showed reactivity against the indicated number of Ad types. (C) Distribution of adenovirus NtAbs by titers. (D) NtAb titers to Ad. The dots indicate individual titer values for each volunteer participating in the study. The dotted line corresponds to the detection threshold. (E) Pearson correlation between seroprevalence and the geometric mean value of NtAb titers to adenoviruses.

Overall, we found that 1.98% of samples exhibited neutralizing activity against all 6 Ad types tested, 6.93% demonstrated neutralizing activity against Ad5, and ∼20% had NtAbs against Ad4 and Ad2 (Fig. 2B). One third of the samples (31.68%) exhibited neutralizing activity against Ad3 and 2.97% did not exhibit neutralizing activity against any of the Ad types tested.

**Figure 2.**
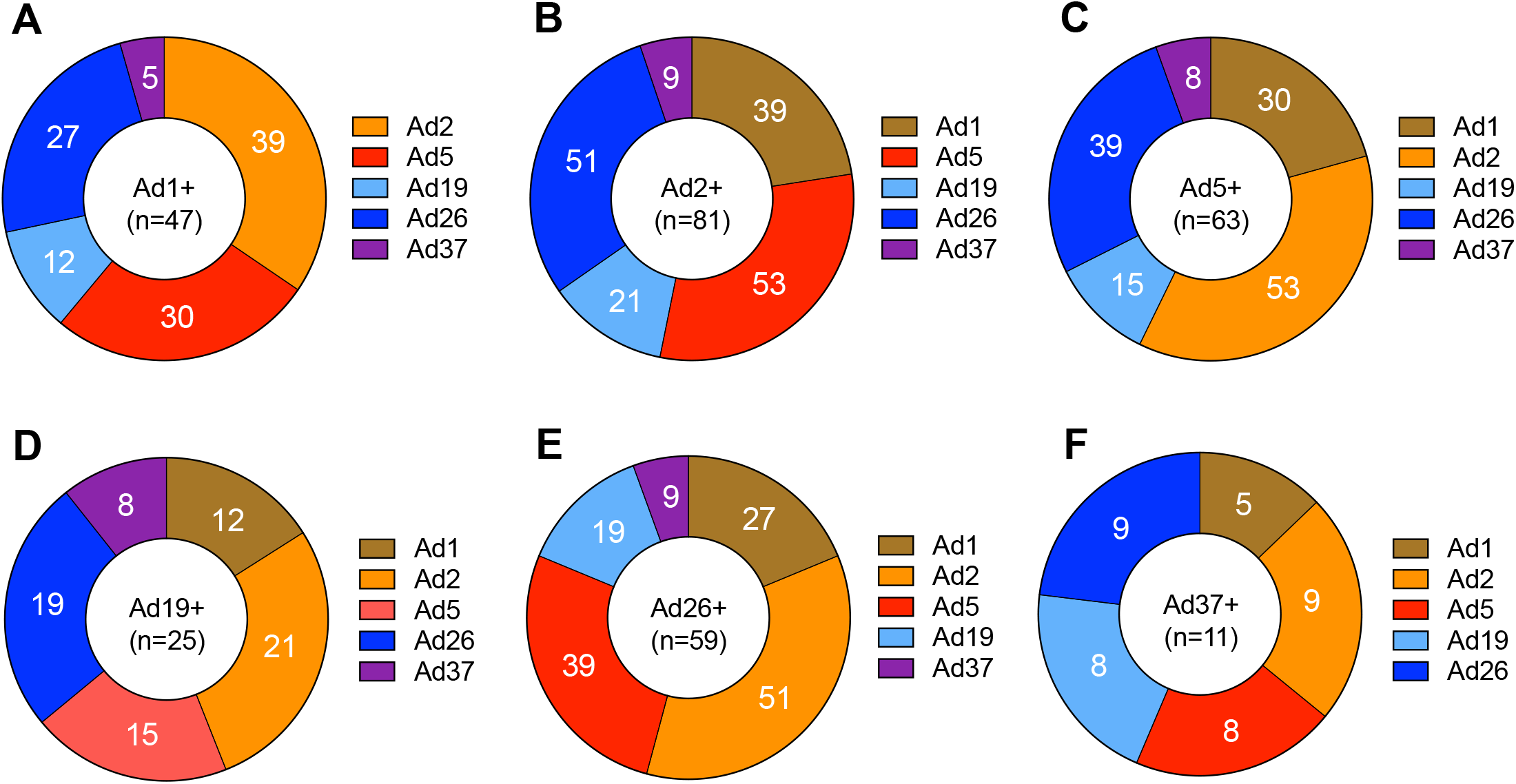
Profiling of the seropositive rates for Ad1 (A), Ad2 (B), Ad5 (C), Ad19 (D), Ad26 (E) and Ad37 (E).

NtAb titers to Ad ranged from <1:25 to 1:12800, which is the minimum and maximum dilution detectable by the assay (Fig. 1D). The distribution plot showed that titers for most volunteers fell within the minimum titer range (≤1:100 – 1:400) for the different adenovirus types (Fig. 1C). The geometric mean NtAb titers to Ad2 were the highest (GMT=266.8) and to Ad1 the lowest (GMT=30.30). The GMTs to Ad19 and Ad37 were below the detection limit (19.13 and 15.04, respectively). The results of statistical differences between the groups are presented in Table 1.

**Table 1.** Comparison of NtAb titers to human adenoviruses.

|  | Ad1 | Ad2 | Ad5 | Ad19 | Ad26 | Ad37 |
| --- | --- | --- | --- | --- | --- | --- |
| Ad1 | - | <0.0001 | 0.0002 | ns | 0.0001 | ns |
| Ad2 | <0.0001 | - | ns | <0.0001 | ns | <0.0001 |
| Ad5 | 0.0002 | ns | - | 0.0001 | ns | <0.0001 |
| Ad19 | ns | <0.0001 | 0.0001 | - | <0.0001 | ns |
| Ad26 | 0.0001 | ns | ns | <0.0001 | - | <0.0001 |
| Ad37 | ns | <0.0001 | <0.0001 | ns | <0.0001 | - |

ns – non-significant

Correlation analysis of the relationship between seroprevalence and the NtAbs GMT to Ad showed a strong direct correlation (r = 0.8313; p = 0.0403) (Fig. 1D). The correlation dependence was exponential: the higher the seroprevalence, the higher the GMT of neutralizing antibodies. We then analyzed in detail the frequency of seropositive samples to different Ad types (Fig. 2). Positive samples to Ad1 (n = 47) were more often positive to Ad2 (n = 39), Ad5 (n = 30) and Ad26 (n = 27). Among Ad2 positive samples (n=81), significantly more were positive for Ad5 (n=53) and Ad26 (n=51) than for Ad1 (n=39) and Ad19 (n=21). Positive samples for Ad5 (n=63) were more often positive for Ad2 (n=53) and Ad26 (n=39) than for Ad19 (n=15) and Ad37 (n=8). Among 59 Ad26 positive samples, NtAbs to Ad2 (n=51) and Ad5 (n=39) were detected most frequently. The number of samples positive to Ad1 was 27, to Ad19 – 19. In samples positive to Ad19 (n=25) or Ad37 (n=11), NtAbs to other types of Ad were detected approximately equally. The smallest number of positive samples was to Ad37.

These findings are likely related to the fact that some people may be susceptible to infection with multiple types of Ad, which cause respiratory infections (Ad1, Ad2, Ad5).

### Cross-neutralization

To investigate whether cross-neutralization is possible between Ad1, Ad2, Ad5, Ad19, Ad26, and Ad37, we studied the correlation between antibody titers in the sera panel. Cross-reactivity of antibodies between Ad1, Ad2, Ad5, Ad19, Ad26, and Ad37 was analyzed using the Spearman’s rank correlation coefficient (Fig. 3). Correlation analysis revealed a significant weak direct association between the NtAb level to Ad19 and to Ad26 (r = 0.2320; p = 0.0196) (Fig. 3M) and a significant moderate direct association between the NtAb level to Ad19 and to Ad37 (r = 0.4401; p < 0.0001) (Fig. 3N). These data indicate that the absence of NtAbs to Ad19 was also accompanied by the absence of NtAbs to Ad26 or Ad37. Most samples did not have NtAbs to Ad19 and Ad37. At the same time, no correlation was found between the NtAb level to Ad26 and Ad37. There was a complete lack of any correlation between other types of adenoviruses, which indicates that the presence of a high titer of antibodies to one type does not lead to the appearance of cross-neutralizing antibodies to another type.

**Figure 3.**
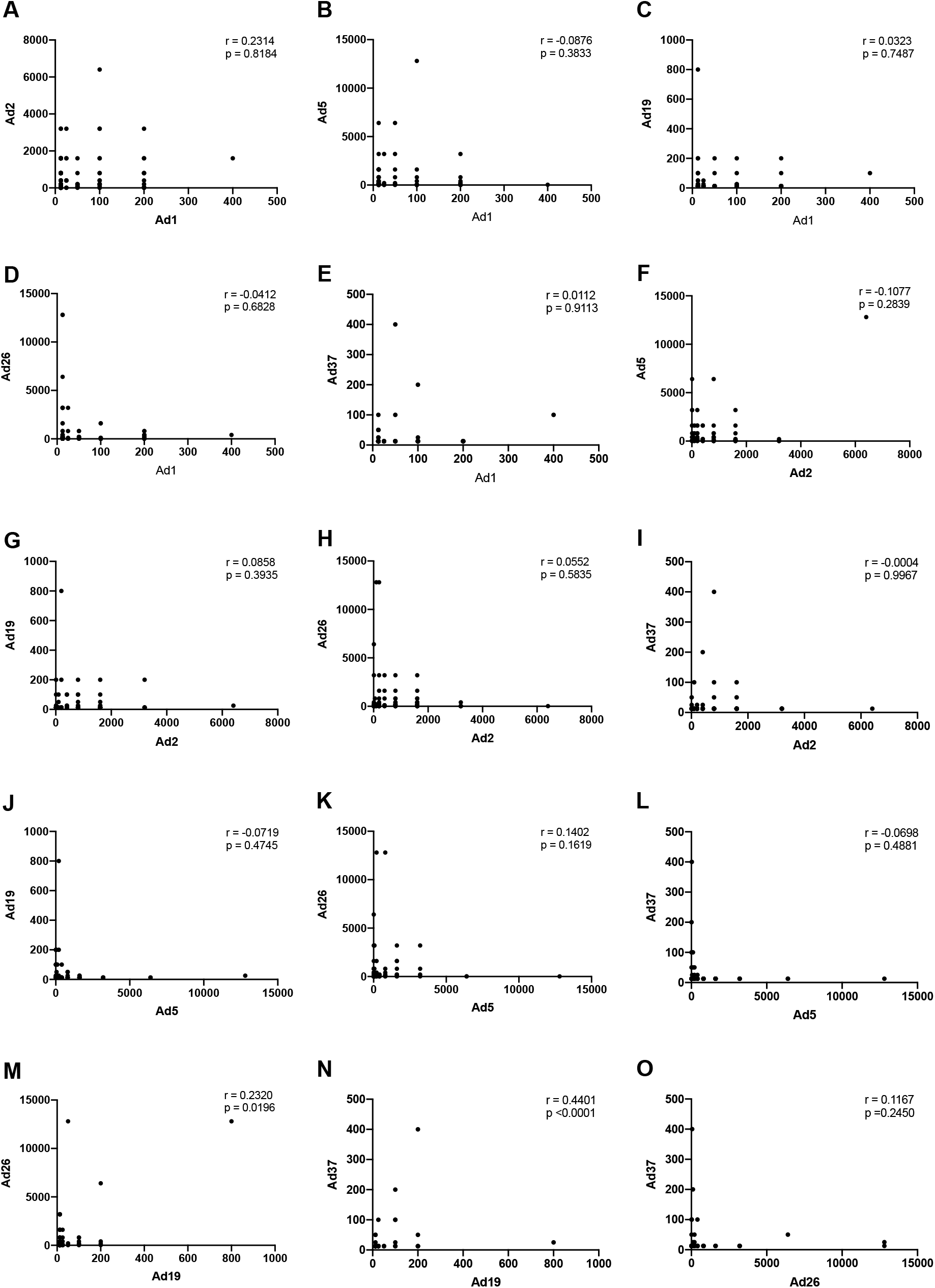
Cross-reactivity of neutralizing antibodies to Ad. Correlation between NtAb titers to Ad5 measured in the same panel of sera. Statistical evaluation was performed using Spearman’s rank correlation coefficient.

## Discussion

The effective use of adenoviral vectors for vaccine development is based on the ability to rapidly induce protective immune responses that persist for a long time. The exceptional immunogenicity of Ad5-based vectors is due to a number of properties: (i) high levels of transgene expression, (ii) activation of innate immunity, (iii) transduction of immature dendritic cells leading to their maturation into antigen-presenting cells, and (iv) a sufficiently long time of antigen presentation due to the lack of induction of apoptosis of transduced cells.

Pre-existing natural immunity to Ad5 is a limitation to the frequent use of Ad5-based vectors in gene therapy and vaccination. To address this issue, increasing the dose may be an option, but this approach would be limited by potential issues with drug toxicity and significant complication of the manufacturing process. Therefore, the primary objective in developing an effective adenovirus vector vaccine will be to explore the possibility of using alternative vectors with lower seroprevalence than Ad5 but equivalent immunogenicity. Several strategies could be chosen to address this issue, the main ones being the use of rare human adenovirus types and the use of adenoviruses from other animal species [15, 37-39]. However, the widespread use of adenoviruses from other animal species is limited not only by production technologies, but also by insufficient knowledge of the association of such adenoviruses with human diseases - these are parameters that require further careful study. Therefore, the use of rare human adenovirus types as potential alternatives to Ad5 is currently a more attractive strategy.

The data presented in this manuscript demonstrate the prevalence of neutralizing antibodies to different types of adenoviruses in healthy individuals. Although we found high NtAb levels to some Ad types, we also showed very low reactivity to other types, making them potential candidates for the development of adenovirus-based vectors in the future.

To determine NtAbs to adenoviruses, different research groups use different methods: methods based on CPE [12], green fluorescent protein [40], luciferase [23], and β-galactosidase reporter virus [41]. The use of different methods, different threshold titers, and other parameters make it very difficult to compare the results of different studies.

From a technical point of view, it should be noted that all types of Ad used in this study were purified by the same cesium chloride gradient ultracentrifugation method and had similar values for the number of viral and infectious particles (Table 2). The existing degree of variability in the ratio of viral to infectious particles did not affect the reliability of the results, as confirmed by the use of an internal positive control sample.

**Table 2.** Characteristics of Ad types and conditions of neutralization reaction.

| Ad type | Reporter gene | Purification method | Concentration |  | Cell line | Incubation time (h) |
| --- | --- | --- | --- | --- | --- | --- |
|  |  |  | vp/ml (OD<sub>260</sub>) | TCID<sub>50</sub>/ml |  |  |
| 1 | EGFP | CsCl | $1.14 \times 10^{12}$ | $2.15 \times 10^9$ | HEK293 | 48 |
| 2 | EGFP | CsCl | $1.59 \times 10^{12}$ | $1.00 \times 10^{10}$ | HEK293 | 48 |
| 5 | EGFP | CsCl | $3.03 \times 10^{12}$ | $6.31 \times 10^{10}$ | HEK293 | 48 |
| 19 | EGFP | CsCl | $2.42 \times 10^{12}$ | $1.31 \times 10^9$ | HEK293 | 48 |
| 26 | EGFP | CsCl | $4.24 \times 10^{12}$ | $1.58 \times 10^9$ | HEK293 | 72 |
| 37 | EGFP | CsCl | $1.46 \times 10^{12}$ | $1.00 \times 10^9$ | HEK293 | 72 |

It is important to note that the seroprevalence of neutralizing antibodies to many types, including Ad5, in our cohort was not as high as previously reported. In the Vogels et al. study [15], the highest proportion of seropositive samples was found for Ad5 (∼80%) and Ad1 (∼70%). For Ad2, this proportion was approximately ∼65%. This is in contrast to our data: while we also saw a high frequency of serum samples with neutralizing activity to Ad5, we saw an even higher reactivity to Ad2, but significantly lower reactivity to Ad1, to which only 46.53% of samples in our cohort showed neutralizing activity. With regard to Ad2, our data are consistent with the results of Wang X. et al. and Klann P.J. et al. studies by [14-16], but differ for Ad5 and Ad1. However, the detected antibody titers should not have a negative impact on the use of vectors based on Ad2, for example. In clinical studies, it was shown that antibody titers to Ad5 below 200 do not affect or have a limited effect on the immunogenicity of the vector [42]. Our findings on Ad26 seroprevalence differ from those reported in the studies mentioned above. We found 58.42% seropositive samples, while Vogels et al. and Wang X. et al. found less than 10% [14]. In the Yi et al. study the seroprevalence rate to Ad26 was 47.0% [43]. It can be assumed that the higher percentage of Ad26 seropositive individuals in our study is associated with the use of the Sputnik V vaccine, in which Ad26 is the first component [29].

It should be noted that our data, of course, do not reflect global seroprevalence. It has been repeatedly shown previously that seroprevalence can be much higher in different countries, for example in Africa and Asia compared to Europe and the United States [20, 22, 44]. It may also differ within a country, as was shown for Ad5 in different regions of China [31].

The conducted correlation analysis showed a strong relationship between seropositivity and the geometric mean titer of neutralizing antibodies to adenovirus. An increase in antibody titers indicates the formation of a humoral immune response and population immunity.

Comparing the titers of neutralizing antibodies to Ad1, Ad2, Ad5, Ad19, Ad26 and Ad37 types, we determined that there is no cross-reactivity between NtAbs to Ad1, Ad2, Ad5 and Ad26. In addition to seroprevalence and the absence of cross-reactivity, an important factor to consider when choosing individual Ad types for the development of candidate vaccines is natural immunogenicity. It also affects the strength of transgene-specific immune responses.

In conclusion, we note that based on the presented data, we assume that Ad1 or Ad2 should be considered the most promising candidates for the development of vaccine vectors.

## Author Contributions

Conceptualization, O.V.Z. and D.Y.L.; Formal analysis, O.V.Z., I.V.D and S.D.V.; Funding acquisition, A.L.G.; Methodology, O.V.Z., T.A.O., O.P., I.V.V., D.I.Z., P.P.G. and D.V.V.; Project administration, O.V.Z. and D.V.S.; Supervision, O.V.Z. and D.Y.L.; Writing—original draft, O.V.Z; Writing—review & editing, O.T.A., I.V.D., G.D.M., D.V.S. and D.Y.L.

## Informed Consent Statement

This study was approved by the Biomedical Ethics Committee of the Gamaleya National Research Center for Epidemiology and Microbiology of the Russian Ministry of Health (Protocol No. 81 dated 05/11/2024).

## Data Availability Statement

The data presented in this study are available on request from the corresponding author.

